# Host developmental stage is associated with shifts in the exosphere microbiome of urban-farmed Asian green leafy vegetables

**DOI:** 10.1101/604819

**Authors:** Shruti Pavagadhi, Aditya Bandla, Miko Poh Chin Hong, Shivshankar Umashankar, Yeap Yoon Ting, Sanjay Swarup

## Abstract

Green leafy vegetables (GLV’s) comprise a significant part of urban South-East Asian (SEA) diets and are intensively farmed in SEA cities, including Singapore. Urban farming practices and urban-adapted vegetable cultivars likely select for specific above- and below-ground microbial communities – microbiomes – that assemble in close proximity to the plant tissues – the exosphere. A healthy exosphere microbiome is important for plant growth and safe human consumption. Using 16S rDNA gene amplicon sequencing compositional analyses, we show here that the exosphere microbiome of two commonly-consumed GLV’s – Choy Sum (*Brassica oleracea Alboglabra Group*) and Gai Lan (*Brassica chinensis var. parachinensis*) – dominated by *Gammaproteobacteria, Alphaproteobacteria, Bacteroidia* and *Actinobacteria*. Shifts in exosphere microbiome composition were strongly associated with plant developmental stage. Finally, microbial taxa consistently detected in the exosphere comprise a small subset, which are predicted to harbour plant-beneficial traits.

**Significance:** Among plant crops, GLVs form an integral part of the Asian diet, especially so in Southeast Asia. Some of these GLVs have short life-cycles (∽30-45 days), which makes them suitable for urban farms in terms of cost advantage as short cycle crops are preferred in urban farms. From a food-security perspective, GLVs forms an important target food group and efforts are being made to increase its productivity to meet the increasing food demands. Current farming practices often place lot of importance on chemical fertilizers and nutrient inputs to improve the fertility of non-arable urban lands to increase the crop productivity. Furthermore, farms in urban settings are also associated with anthropogenic inputs and eutrophic conditions. These together, contribute to negative environmental externalities questioning the sustainability and eco-sustenance of urban farming. Microbial based management systems can not only resolve these challenging issues, but can also enhance plant growth, nutrient use efficiency and disease tolerance. However, their use as microbial adjuncts to agricultural practices is currently limited in urban environments, which could possibly be due to the restricted knowledge-base on these urban phytobiomes.

## Observations

### Host developmental stage and host identity shape the exosphere microbiome

Replicated samples (n=3) of two GLV’s – Choy Sum (*Brassica oleracea Alboglabra Group*) and Gai Lan (*Brassica chinensis var. parachinensis*) – across two major plant developmental stages (seedlings and adults) were randomly collected from the best performing greenhouses in one of Singapore’s largest commercial farms involved in green leafy vegetable production. Microbial communities associated with the rhizosphere and phyllosphere – the exosphere, were analysed using 16S rDNA gene amplicon sequencing. A total of 1, 355, 621 (median read count per sample: 25, 775; range: 101 – 41425 reads per sample) high-quality reads were obtained, which in turn, mapped to a total of 12, 735 amplicon sequence variants (ASV’s). Sample counts were total sum scaled and square-root transformed prior to beta-diversity analysis.

Similarity of samples in terms of microbiome composition were visualised using unconstrained principle coordinate analysis (PCoA). Samples separated according to host developmental stage along the first axis for both the rhizosphere and phyllosphere, while separation along the second axis largely corresponded to host identity of adult plants (Figure 1). These patterns were corroborated using Permutational Multivariate Analysis of Variance (PERMANOVA) which showed that host developmental stage accounts for the largest amount of variance in terms of microbiome composition (*SI Appendix, Dataset S1*).

**Figure 1.**
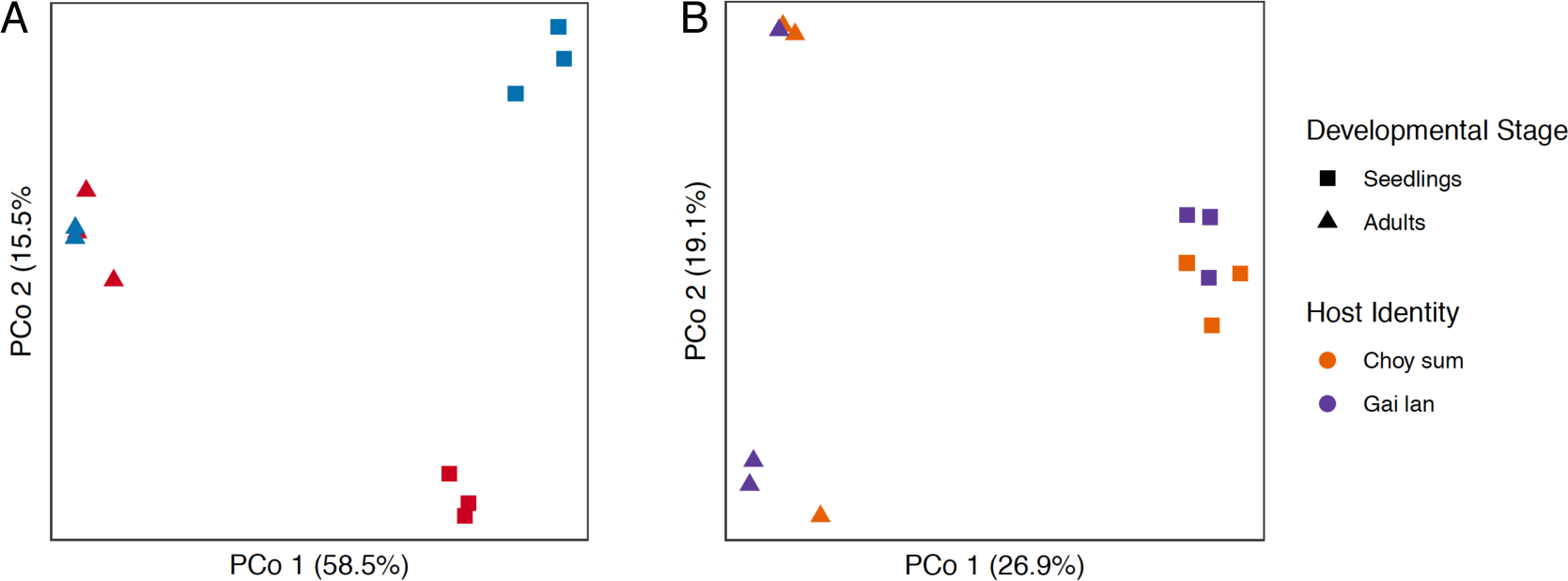
Microbiome composition of the plant exosphere is associated with host developmental stage and host identity. (A) Rhizosphere and (B) Phyllosphere

### Microbial taxa in the exosphere are predicted to harbour plant-beneficial traits

Stable and significantly enriched components of the rhizosphere microbiome were identified through differential abundance analyses as well as retaining only those taxa that were detected across all replicates. We found that only a small subset of ASV’s were significantly enriched in the rhizosphere of seedling and adult plants (*SI Appendix, Dataset S2-S5*). Cumulatively, such ASV’s accounted for 15% and 33% of total microbial community in Choy Sum (1753 ASVs; 36 enriched ASVs) and Gai Lan (1638 ASVs; 81 enriched ASVs) seedlings respectively. While for adult plant types, it accounted for 18.6% and 16.1% of the total microbial community in Choy Sum (3802 ASVs; 219 enriched ASVs) and Gai Lan adults (3233 ASVs; 119 enriched ASVs) respectively.

Stable components of the phyllosphere microbiome were identified as those taxa that were consistently detected across all replicates derived from the respective groups (*SI Appendix, Dataset S6*). Cumulatively, such microbial taxa accounted for 39.4 % and 78.4 % of total microbial community in Choy Sum and Gai Lan seedlings respectively. While for adult plant types, they accounted for 48.7 % and 14.8 % of the total microbial community in Choy Sum and Gai Lan adults respectively. Interestingly, most of these prevalent taxa derive from the phyla *Alphaproteobacteria, Gammaproteobacteria, Deltaproteobacteria, Bacteroidia* and *Actinobacteria*.

Next, we predicted the genomic repertoire of these taxa using PICRUSt2 (Eddy 1998; Langille et al., 2013; Louca and Doebeli, 2018; Barbera et al., 2019, Czech et al., 2019) and then searched for genomic features related to plant beneficial functions (*SI, Appendix, Dataset S7*). Some of these functions pertain to genes involved in nitrogen, sulfur and phosphorus metabolism. These are essential elements for growth, development, and various physiological functions in plants. Furthermore, members from *Brassicaceae* have higher nitrogen, phosphorus and sulfur requirements than other crop species, and therefore, are particularly sensitive to their availability (Walker and Booth, 2003; Albert et al., 2012, Brennan and Bolland, 2009).

We found that genes predicted to be involved in ureolysis were the most prevalent across all sample groups i.e. seedling and adult exospheres of both GLV’s (*SI, Appendix, Dataset S8*). Although, such genes are involved in the hydrolysis of urea to ammonia, we found ammonia oxidizers to be highly abundant both in the bulk soil and the rhizosphere. We speculate that these taxa may act synergistically to convert urea to plant available nitrates. These nitrates form an integral component of many structural and metabolic compounds in plant cells such as amino acids, proteins, nucleotides, chlorophyll, chromosomes, genes and all enzymes. Nitrogen deficiency in some members from *Brassicaceae* is associated with lowered sensitivity to water stress (Albert et al., 2012)

Predicted genomic features associated with phosphate solubilization and sulfate oxidation were also consistently detected in multiple taxa (*SI, Appendix, Dataset S8*). Both these nutrients are limited in nature and they are often present in unavailable chemical forms in the soil. Sulfate and phosphate oxidizing microbes convert these to more readily available substrates that can be utilized by the plants. Both, phosphorus and sulfur are essential nutrients required for formation of structural components in plants, including nucleic acids, phospholipids and a variety of secondary metabolites, which play a pivotal role in protecting these plants against abiotic and biotic stressors (Plaxton and Lambers, 2015).

## Conclusions

These results show that host developmental stage is associated with major shifts in the exosphere microbiome of widely-consumed urban-farmed Asian GLVs. Further, taxa that were consistently detected in the exosphere are predicted to harbour plant-beneficial functions.

## Supporting Information (SI) Appendix

SI Dataset 1: PERMANOVA results for rhizosphere and phyllosphere microbiome

SI Dataset 2: ASVs significantly enriched in the seedling choy sum rhizosphere

SI Dataset 3: ASVs significantly enriched in the adult choy sum rhizosphere

SI Dataset 4: ASVs significantly enriched in the seedling gai lan rhizosphere

SI Dataset 5: ASVs significantly enriched in the adult gai lan rhizosphere

SI Dataset 6: ASVs consistently detected in all replicates of respective phyllosphere

SI Dataset 7: List of plant-beneficial functional genes

SI Dataset 8: Frequency of predicted plant beneficial functions in exosphere microbiota

## Acknowledgements

This work was supported by the National Research Foundation, Prime Minister’s Office, Singapore under its Competitive Research Programme (NRF-CRP16-2015-04). We also thank the NUS Environmental Research Institute (NERI) and Singapore Centre for Environmental Life Sciences Engineering (SCELSE) for their services and support.

## Author’s contributions

S.P., S.U. and S.S. conceived the study. S.P. and M. P. C. H. were involved in sampling and sample preparation. A.B. was involved in data analysis. A.B. and S.P. were involved in data interpretation and writing the manuscript.

## Statement of informed consent, human/animal rights

No conflicts, informed consent, human or animal rights applicable.

